# Somatostatin-excitatory coordination emerges with dexterous motor skill learning

**DOI:** 10.64898/2026.10.28.762594

**Authors:** Gabriella Wheeler Fox, Eliza T. Wiener, Jason N. MacLean

## Abstract

Motor skill-learning increases spatio-temporal consistency in neuronal activity in primary motor cortex (M1), and Somatostatin-expressing (SST) interneurons are causally involved in these learning-related activity changes. Yet, how SST-excitatory interactions shape population level activity changes during motor skill acquisition is still unknown. Here, we imaged large populations of excitatory neurons and identified SST or parvalbumin-expressing (PV) interneurons across successive days as mice learned a dexterous, persistently M1-dependent reach-to-grasp task. Sparse, temporally coordinated, trial-resolved, and rate-controlled functional interactions between SST neurons and skill-modulated excitatory neurons emerged with learning and were stronger than those in excitatory-excitatory or PV-excitatory pairs. Interactions strengthened as neurons were recruited to the skill-modulated ensemble and weakened when skill modulation was lost. SST activity also became temporally reliable earlier than excitatory or PV activity, and SST consistency was significantly associated with behavioral performance even after controlling for training session number. Additionally, SST neurons carried more information about movement kinematics than randomly selected subsets of excitatory neurons. Overall, SST neurons became selectively coordinated with behaviorally relevant excitatory populations, revealing a cell-type-specific functional architecture for dexterous motor learning.

## Introduction

Primary motor cortex (M1) undergoes substantial functional and structural reorganization during motor skill learning. As movements become more skilled, initially variable patterns of cortical activity become increasingly reproducible and temporally structured, accompanied by learning-related remodeling of excitatory synapses [1][2]. Inhibitory neurons are active parts of this reorganization. Somatostatin-expressing (SST) interneurons regulate learning-dependent synaptic remodeling and are linked to the emergence of structured pyramidal-cell activity. Parvalbumin-expressing (PV) interneurons undergo complementary plasticity that may contribute to stabilization of evolving cortical activity [3, 4]. Yet how SST and PV interneurons become functionally coordinated with excitatory neurons as motor skill related activity patterns emerge is poorly understood.

Motor learning produces cell-type specific remodeling of inhibitory circuits. Results thus far have supported a model in which learning is facilitated by reduced SST-mediated inhibition. SST axonal boutons are reduced, whereas PV boutons transiently increase [3], and sensory associative learning is accompanied by reduced SST synaptic strength [5] and decreased correlation of SST neurons with the local network [6]. In contrast, PV interneurons become increasingly selective for task-relevant stimuli and more strongly coupled to excitatory neurons with similar selectivity [6]. However, several findings argue against a simple global withdrawal of SST influence. SST population activity increases during motor learning over the same period in which SST boutons are lost [7], and either increasing or decreasing SST activity during training disrupts learning and alters spine stability [3]. Moreover, SST interneurons are causally required for learning-related changes in the timing of M1 pyramidal-cell activity [4]. Together, these findings suggest that the contribution of SST inhibition depends not simply on its overall magnitude, but on which neurons are engaged and when. Resolving this apparent dissociation between reduced structural or synaptic inhibition and increased SST recruitment therefore requires determining how SST neurons are functionally coordinated with excitatory populations during learning, and how this organization differs from that of PV interneurons.

The organization of inhibitory activity may also depend strongly on the demands that a learned behavior places on M1. Motor cortex can become dispensable for execution of some extensively trained lever-press behaviors [8, 9], and SST recruitment itself varies with task demands: during a single-lever-press task, deep-layer SST neurons are strongly action locked only during acquisition, whereas action locking persists longer during acquisition of a more temporally constrained lever-press sequence [10]. It remains unclear how SST and PV interneurons become coordinated with excitatory populations during acquisition of dexterous skills that continue to require M1 after learning is established.

We address these questions by analyzing cell-type specific functional interactions during acquisition of a single-pellet reach-to-grasp skill that requires continuous, online control of movement [11] and remains dependent on M1 even after prolonged training [12–16]. We asked whether SST and PV interneurons become selectively coordinated with particular excitatory neurons during learning, and whether the excitatory neurons participating in these interactions are especially relevant to kinematic decoding. We imaged large populations of M1 neurons across successive days of training, using static fluorescent labels to identify SST interneurons in one cohort and PV interneurons in another. To quantify interactions across cell types, while controlling for differences in activity rate, we developed a mutual-information analysis combining adaptive discretization with a rate-preserving temporal shuffle.

This analysis revealed a sparse and highly specific organization of SST-excitatory interactions that emerged during skill learning. SST interneurons became strongly coordinated with a restricted subset of skill-modulated excitatory neurons, in contrast to the broad decorrelation reported for SST neurons during visual associative learning. PV neurons were initially strongly coordinated with skill-modulated excitatory neurons, but became increasingly decoupled from the skill-modulated network with learning. Temporal consistency in SST activity emerged earlier during learning and ultimately exceeded that of excitatory or PV neurons, with early SST consistency correlating with improvements in behavioral performance. Moreover, both SST neurons and excitatory neurons strongly functionally coupled to SST neurons carried more information about movement kinematics than randomly selected subsets of excitatory neurons. Together, these results identify selective SST-excitatory coordination as a prominent feature of the functional architecture that develops during acquisition of a dexterous, M1-dependent skill. Rather than simply disengaging to permit plasticity, SST neurons become selectively and temporally coordinated with a behaviorally relevant excitatory ensemble, while PV neurons are initially highly coordinated with excitatory neurons and then become decoupled. These findings suggest that SST inhibition is organized with cellular and temporal specificity and that SST and PV interneurons make distinct contributions according to the demands placed on the circuit.

## Results

### SST activity becomes increasingly consistent during acquisition of a dexterous reach-grasp-carry skill

We trained mice to reach for, grasp, and carry a sucrose pellet from a pedestal to the mouth using the right forelimb (fig 1a). This skill requires complex paw and digit kinematics and remains dependent on motor cortex for performance, not just learning [12–16]. Following initial shaping, during which pellets were presented in a trough, the pedestal was placed at a fixed position optimized for each mouse. Performance improved rapidly over the first several days of training: success rate reached a plateau by approximately day 3 (fig 1b), whereas the number of successful pellet retrievals per session continued to increase until approximately day 7 (fig 1c). Since the task was self-paced, the continued improvement in success count indicates improved efficiency.

**Fig. 1:**
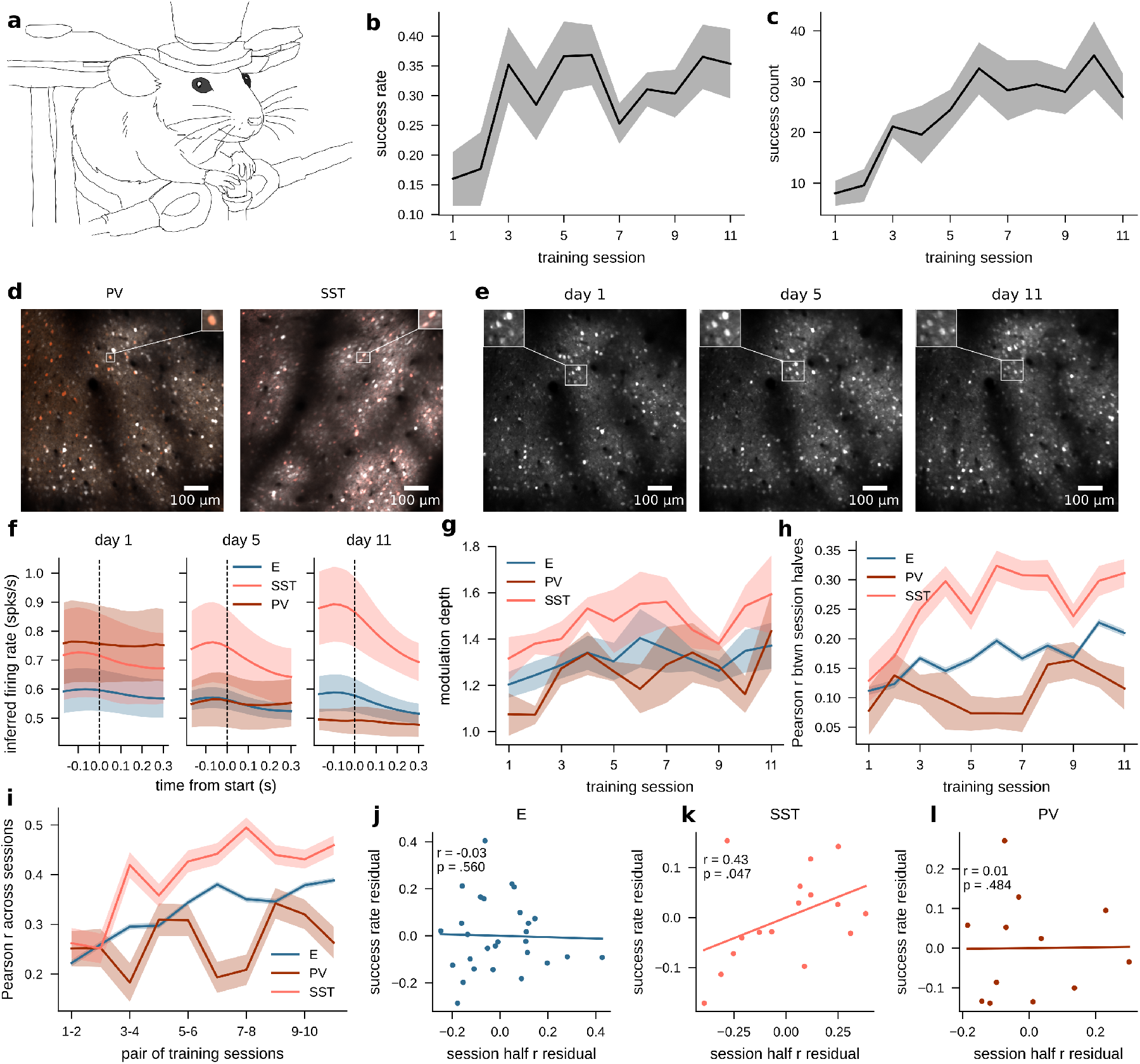
SST activity becomes increasingly consistent during acquisition of a dexterous reach-grasp-carry skill. **a.** Schematic of the pellet reach task setup. **b.** Success rate over training sessions. Shading shows SEM over mice (n=7). Logistic regression per mouse on binary trial outcomes vs session number for first 3 sessions, with Wald test on coefficients: mouse 1: coef = 0.781, p=0.002, CI [0.284 1.27]; mouse 2: coef = 0.626, p=0.007, CI [0.174 1.079]; mouse 3: coef = 0.574, p = 0.008, CI [0.151 0.996]; mouse 4: coef = 0.655, p = 0.024, CI [0.088 1.222]; mouse 5: coef = -0.126, p = 0.597, CI [-0.593 0.341]; mouse 6: coef = 0.906, p = 0.006, CI [0.256 1.556]; mouse 7: coef = 1.57, p = 0.000, CI [0.831 2.316]. **c.** Success count over sessions. Shading shows SEM over mice (n=7). Linear mixed effects model with mice as random effect: on sessions 1–3, coef = 6.571, p = 0.000, CI 3.092 to 10.05; on days 4–11, coef = 0.449, p = 0.473, CI -0.776 to 1.674. **d.** Mean GCamp6f fluorescence overlaid with tdTomato labels for PV (left) and SST (right) neurons. **e.** Field of view for an example mouse on sessions 1, 5, and 11. **f.** Mean population activity by cell type with SEM over mice (n=7) for sessions 1, 5, and 11. **g.** Modulation depth relative to the median of activity during non-skill epochs over sessions with SEM over mice (n=7). **h.** Correlation between PETHs for randomly split halves of sessions, with 1000 random draws. SEM over neurons per session (n = 740.8 *±* 213.4 SD E neurons, n = 76.5 *±* 27.8 SD SST neurons, n = 44.9 *±* 23.0 SD PV neurons). Linear mixed effect model for correlation versus session pair, with mice as a random effect and number of trials an additional covariate. Days 1–3: E: coef = 0.025, p = 0.132, CI [-0.007 0.057], SST: coef = 0.061, p = 0.038, CI [0.003, 0.118], PV: coef = -0.006, p = 0.75, CI [-0.041 0.029]; Days 1–11: E: coef = 0.006, p =0.003, CI [0.002 0.009]; SST: coef = 0.011, p = 0.003, CI [0.004 0.018]; PV: coef = 0.002, p = 0.509, CI [-0.004 0.009]. **i.** Pearson correlation between PETHs across sessions for longitudinally tracked neurons, with SEM over neurons per adjacent session pair (n=514 *±* 188.6 SD tracked E neurons, n = 59.3 *±* 24.8 SD tracked SST neurons, n = 31.5 *±* 18.5 SD tracked PV neurons per adjacent session pair). Linear mixed effect model for correlation versus session pair, with mice as a random effect and number of trials as an additional covariate. Days 1–3: E: coef = 0.04, p = 0.033, CI [0.003 0.076]; SST: coef = 0.136, p = 0.002, [0.049 0.224]; PV: coef = -0.04, p = 0.066, CI [-0.083 0.003]. Days 1–11: E: coef = 0.013, p = 0.000, CI [0.007 0.018], SST: coef = 0.022, p = 0.012, CI [0.012 0.032], PV: coef = 0.000, p = 0.988, CI [0.000 0.001]. **j–l.** Partial correlation between session half correlation and success rate, controlling for session number, on first four sessions per mouse. Pearson r and Wald test vs 0: j. n = 28 sessions, r=0.03, p = 0.56 k. n = 16 sessions, r=0.43, p = 0.047 l. n = 12 sessions, r = 0.01, p = 0.48.

On each training day, we performed two-photon calcium imaging of neurons expressing GCaMP6f under a pan-neuronal promoter (803.7 *±* 233.5 neurons per session) in the caudal forelimb area (CFA) of left hemisphere primary motor cortex. SST interneurons (76.5 *±* 27.8 neurons per session; n=4 mice) and PV interneurons (44.9 *±* 23 neurons per session; n = 3 mice) were identified with tdTomato labeling (fig 1d). For each recording session, we aligned our imaging field of view with a reference frame from the previous day, allowing us to identify the same neurons across sessions using CellReg (fig 1e).

We first examined how the temporal consistency of skill-related neural activity changed during learning. We identified individual reach, grasp, and carry movements within the self-paced behavioral sequence. Because mice frequently executed multiple reaches before grasping an object and multiple grasps before initiating a carry, these movements did not occur in a fixed one-to-one sequence. We therefore analyzed each movement type separately, aligning neural activity to the onset of individual reach, grasp, or carry events. Across training, SST neurons exhibited increased skill-related activity and greater temporal consistency of movement-aligned responses than either putative excitatory or PV neurons (Fig. 1f). Modulation depth, quantified as the peak population activity during the skill divided by the median of non-skill population activity, was also highest for SST neurons and increased with learning (fig 1g).

At the single cell level, SST neurons also showed increasingly consistent temporal activity patterns. Within-session consistency, quantified as the correlation between peri-event time histograms (PETHs) computed from randomly selected session halves including all reach, grasp, and carry trials, increased across training (fig 1h). Across-session consistency, quantified as the correlation between PETHs on adjacent training days for longitudinally matched neurons, showed a similar increase (fig 1i). These analyses included all recorded neurons rather than selecting cells based on skill modulation or trial-to-trial reliability. By the third day of training, SST neurons were more consistent than both putative excitatory and PV neurons, and this difference persisted on subsequent days. PV neurons, in contrast, exhibited the lowest consistency both within and across sessions.

We next asked whether variation in neural consistency across sessions was related to behavioral performance. Because both success rate and PETH consistency increased with training day, their raw correlation could arise simply from this shared temporal trend. We therefore calculated the partial correlation between PETH consistency and behavioral performance while controlling for training day. SST consistency remained significantly associated with performance after accounting for session number, indicating that sessions with more reliable SST activity were associated with better performance than expected from learning progression alone (fig 1k, *r* = 0.43*, p* = 0.047). PV and putative excitatory PETH consistency was not related to success rate (figs 1j,l). Thus, as SST neurons became increasingly recruited during skilled behavior, their activity also became more temporally reliable, and session-to-session variation in this reliability tracked behavioral performance beyond the overall effects of training.

### A skill-modulated network emerges during skill learning

We next asked how the identities of neurons involved in the skill changed across learning. For each neuron, we tested whether its activity differed between the skill-related epochs and other timepoints using an ANOVA, with Bonferroni correction for the number of neurons tested per session. The fraction of skill-modulated neurons increased over the first few days of training across all cell types, indicating recruitment of neurons into the task-related activity pattern (fig 2a). The fraction of skill-modulated neurons was significantly greater for SST and PV interneurons than for excitatory neurons, with this difference becoming more pronounced with learning. Most skill-modulated neurons showed increased, rather than decreased, activity during the skill epochs; we therefore refer to these neurons as the skill-modulated network in subsequent analyses. Fig 2b shows example trial averaged activity for skill-modulated neurons. We next asked whether this selective recruitment was accompanied by structured interactions.

**Fig. 2:**
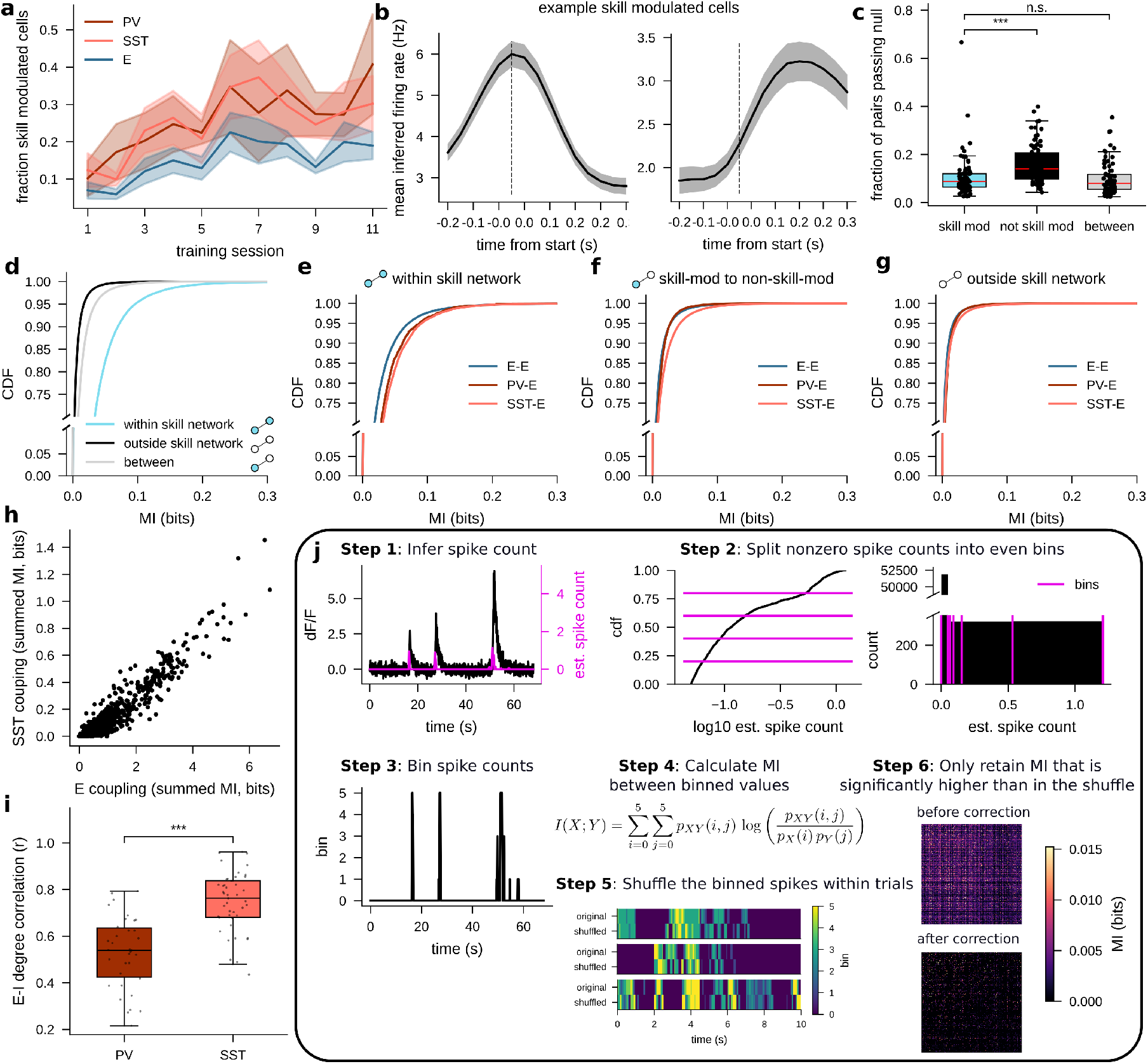
A skill-modulated network emerges during skill learning. **a.** Fraction of up-modulated neurons across sessions by cell type. Shading shows SEM over mice (n = 7). Linear mixed effects model with mice as a random effect. Sessions 1–6: putative excitatory: coef = 0.029, p = 0.000, CI [0.017 0.041]; SST: coef = 0.042, p = 0.004, CI [0.014, 0.070]; PV: coef = 0.041, p = 0.016, CI [0.008 0.074]. The fraction of up-modulated neurons is significantly greater for PV and SST neurons after session three, but not sessions 1-2. Mann Whitney U-tests sessions 1-2: E vs PV: p = 0.13; E vs SST p = 0.095. Sessions 3-11: E vs PV: p = 0.0014; E vs SST: p = 0.0003. **b.** Example trial-averaged inferred firing rates from two up-modulated neurons for an example mouse. Dashed line shows trial start. **c.** Fraction of pairs that pass the null in the skill-modulated network (cyan), in non-network pairs (black), and pairs between skill-modulated and non skill-modulated (gray). N = 79 sessions. *** indicates p¡0.001 on Mann-Whitney U-test. Within vs non skill-modulated rank biserial r = 0.49. **d.** Empirical CDFs of MI within the skill-modulated network (cyan, n = 46950 pairs), between skill-modulated and non-skill modulated neurons (gray, n = 396290 pairs), and between non skill-modulated neurons (black, n = 2547497 pairs). Mann-Whitney U-tests: within vs not skill-modulated: p = 0.0, rank biserial r=0.73; within vs between: p = 0.0, rank biserial r = 0.54; between vs not skill-modulated: p = 0.0, rank biserial r = 0.28. **e.** Empirical CDFs of MI within the skill network separated into SST-excitatory pairs (n = 12208), excitatory-excitatory (n = 76079), and PV-excitatory (n = 2052) pairs. Mann-Whitney U-test: SST-E vs E-E p = 0.0, rank biserial r = 0.22; PV-E vs E-E p = 1.1e-63, rank biserial r = 0.22. **f.** Empirical CDFs of MI between skill-modulated and non skill-modulated neurons separated into SST-excitatory pairs (n = 65583), excitatory-excitatory (n = 544658), and PV-excitatory (n = 19733) pairs. Mann-Whitney U-test: SST-E vs E-E p = 0.0, rank biserial r = 0.11; PV-E vs E-E p = 7.8e-228, rank biserial r = 0.14. **g**. Empirical CDF of MI in non skill-modulated pairs, separated into SST-excitatory pairs (n = 297885), excitatory-excitatory (n = 2880709), and PV-excitatory (n = 104148) pairs. Mann-Whitney U-test: SST-E vs E-E p = 0.0, rank biserial r = 0.045; PV-E vs E-E p = 0.0, rank biserial r = 0.21. **h.** Excitatory coupling versus SST coupling for an example session. N = 1015 excitatory neurons in session. **i.** Pearson correlation between E and I coupling for PV (n = 34 sessions for 3 mice) and SST (n = 45 sessions for 4 mice). *** indicates *p <* 0.001 on Mann-Whitney U-test. Rank biserial r = 0.71. **j**. Schematic describing the mutual information calculation procedure, including adaptive binning to discretize activity and comparison against a trial-rate matched shuffle null.

We compared pair-wise interactions within the skill-modulated network, within the non-network population, and between network and non-network neurons. To identify temporally structured interactions while controlling for differences in activity rate, we combined mutual information (MI) with adaptive discretization and a rate-preserving temporal shuffle (fig 2j). Inferred firing rates were dis-cretized using adaptive bins, with nonzero observations divided into bins of equal occupancy across the session. This approach preserved graded structure in the inferred firing rates while accommodating their strongly skewed distribution. For each neuronal pair, we then calculated MI based on the discretized activity states across concatenated trials.

Because skill-modulated and non-skill-modulated neurons differed systematically in their activity rates, raw MI could differ simply because of differences in their marginal activity distributions. We therefore compared the observed MI with a null distribution generated by independently circularly shifting activity within each 450-ms trial by 90–300 ms (3–10 imaging bins) and recalculating MI across 1,000 shuffles. This procedure preserves each neuron’s activity distribution and per trial mean firing rate while disrupting the temporal alignment between neurons. We only retained pairs whose observed MI significantly exceeded the shuffled distribution (14.59 *±* 7.81% of pairs). Thus, the resulting MI identifies pairwise dependence that depends on temporal coordination within individual trials on a sub-second timescale and cannot be explained by differences in activity rate alone. Because calcium signals and inferred firing rates themselves integrate activity over time, this measure should be interpreted as temporally structured coordination between neurons rather than millisecond-scale synchrony or evidence of direct synaptic coupling.

The proportion of significant interactions within the skill-modulated network was lower than among non-modulated neurons (fig 2c), and among pairs with significant interactions, MI was highest between neurons within the skill-modulated network (fig 2d). Thus, interactions between skill-modulated neurons were sparse and comparatively strong, showing that neurons preferentially active during the skill had more reliable trial-to-trial temporal coordination with one another during the skill than other neurons, beyond co-modulation.

This organization was particularly pronounced for inhibitory interneurons. Both SST-excitatory and PV-excitatory pairs had significantly greater MI than excitatory-excitatory pairs within the network (fig 2e). Between skill-modulated and non-skill-modulated neurons, SST-excitatory MI exceed PV-excitatory and excitatory-excitatory (fig 2f). Differences between pairs were absent outside the skill-modulated network (fig 2g). SST and PV neurons were therefore not only preferentially recruited during skilled behavior, but also strongly coordinated with a restricted subset of excitatory neurons within the skill-modulated network.

Elevated MI can arise from aligned activity, in which similar activity states co-occur more often than expected, or from anti-aligned activity, in which elevated activity in one neuron preferentially coincides with low or absent activity in another (similar to a negative correlation). We therefore asked whether the elevated MI observed in PV-excitatory and SST-excitatory pairs arose from aligned or anti-aligned activity. We decomposed MI into its pointwise components (17; see also 18, for a similar method), and grouped subsets of the components into aligned and anti-aligned interactions according to the proximity of the two activity states (see Methods and fig S1a-c for details). Across all cell types, aligned patterns of activation contributed substantially more to total MI than anti-aligned patterns (fig S1d-e). The elevated MI between PV-excitatory and SST-excitatory pairs relative to excitatory-excitatory pairs is also reflected in the aligned patterns (fig S1d), indicating that this MI difference arises from aligned patterns of activity. Thus, the elevated MI between SST and PV interneurons and skill-modulated excitatory neurons was primarily the result of temporally aligned activity rather than preferential anti-correlation of their activity states.

Inhibitory–inhibitory pairs also exhibited high MI (fig S2) regardless of whether they belonged to the skill-modulated network, consistent with prior work showing coordinated activity among inhibitory interneurons, particularly SST cells [19, 20].

We were also interested in the overall coupling structure in the population. If an excitatory neuron was strongly coordinated with SST or PV neurons, how was it positioned in the rest of the network? To test this, we measured the excitatory and inhibitory coupling strength for each excitatory neuron as the sum of its significant MI with other neurons (i.e. weighted degree). SST and excitatory coupling were highly correlated (example session in fig 2h, all sessions in 2i), and PV and excitatory coupling were moderately correlated (2i). This indicates that excitatory neurons highly coupled to SST neurons were more likely to be highly coupled to other excitatory neurons, and vice versa, revealing a tightly integrated set of excitatory neurons that were coordinated with each other and with SST neurons.

### Elevated SST-excitatory coordination emerges with learning

We next asked whether these selective interactions between excitatory and inhibitory neurons changed across learning. Elevated MI within the skill-modulated network was present from the very beginning of training (fig 3a), which was striking considering that the identity of skill-modulated neurons was still highly variable, and single neuron trial averaged activity was inconsistent. Additionally, MI in PV-excitatory and SST-excitatory pairs varied across early and late learning (fig 3b). In the first several sessions, PV-excitatory pairs had the greatest MI, then MI in SST-excitatory pairs increased relative to other cell type pairs, becoming substantially greater around session 7. This increased difference was driven both by reduction in excitatory-excitatory and PV-excitatory MI, as well as increases in SST-excitatory MI. Thus, selective coordination between PV and excitatory cells is present in early sessions, while selective coordination between SST and excitatory neurons emerges later. A substantial literature has measured behavioral and neural differences across acquisition and refinement phases of motor learning [21–26]. Our results here are consistent with the idea that distinct neural changes support distinct phases of learning.

**Fig. 3:**
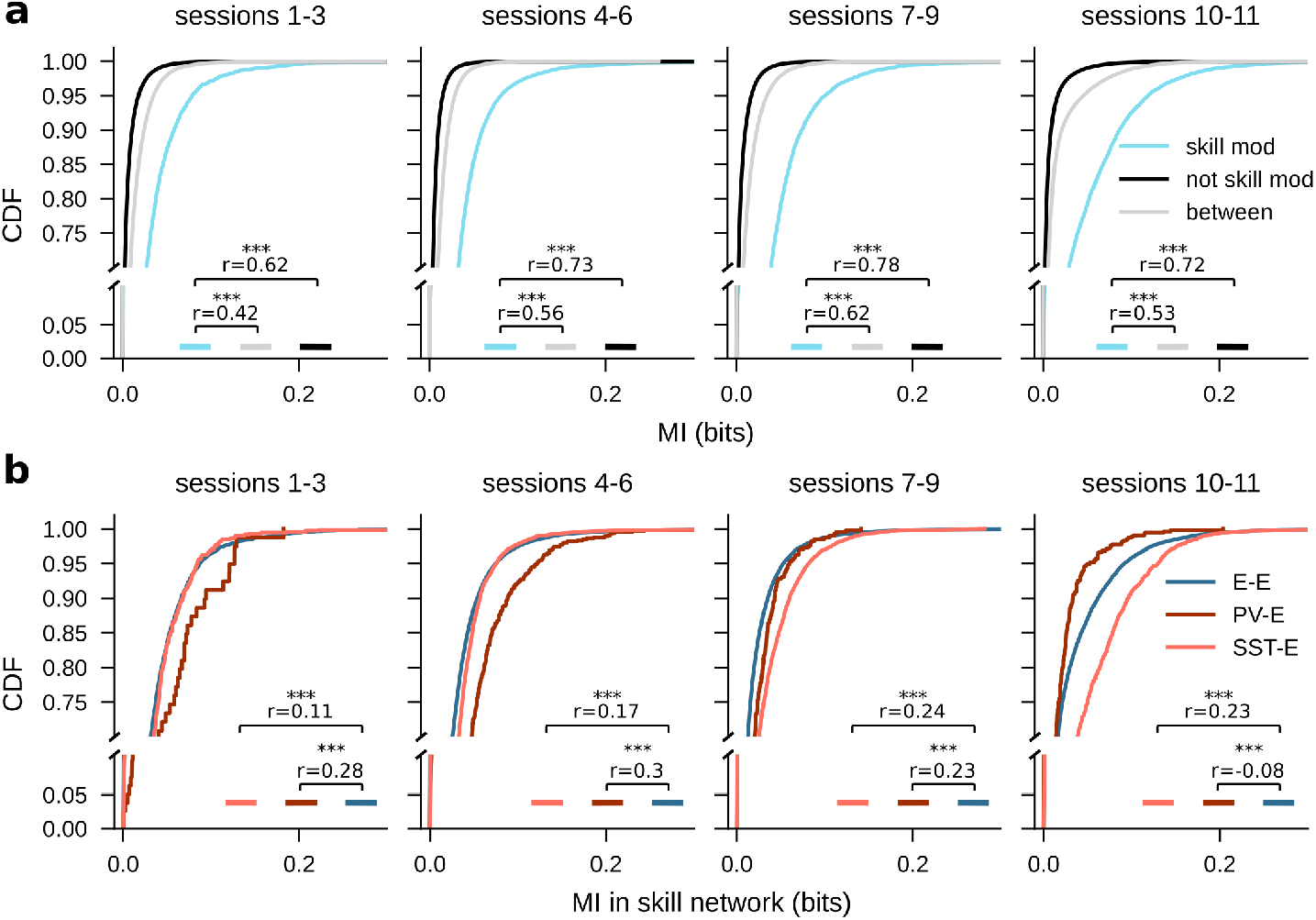
Elevated SST-excitatory coordination emerges with learning. **a.** CDFs of MI within the skill-modulated network, between skill-modulated and non-skill modulated neurons, and in non skill-modulated pairs for sessions 1-3 (n = 4338, 46508, and 572725 pairs), sessions 4-6 (n = 21200, 139375, and 689400 pairs), sessions 7-9 (n = 11527, 118639, and 738059 pairs), and sessions 10-11 (n = 9362, 79678, and 446890 pairs). *** indicates p¡0.001 on Mann-Whitney U-test; r indicates rank biserial correlation. **b.** CDFs of MI within the skill-modulated network separated by cell type and binned into groups of sessions: sessions 1-3 (n = 3117 E-E pairs, n = 487 SST-E pairs, n = 64 PV-E pairs), sessions 4-6 (n = 19153 E-E pairs, n = 3523 SST-E pairs, n = 555 PV-E pairs), sessions 7-9 (n = 26553 E-E pairs, n = 3732 SST-E pairs, n = 343 PV-E pairs), and sessions 10-11 (n = 12149 E-E pairs, n = 1882 SST-E pairs, n = 459 PV-E pairs). *** indicates p¡0.001 on Mann-Whitney U-test, r indicates rank biserial correlation.

### Changes in functional interactions track skill-modulated network recruitment and pruning

We next asked whether changes in network membership were accompanied by corresponding changes in functional interactions. The fraction of skill-modulated neurons newly recruited from one session to the next was highest early in training, and declined with learning, consistent with progressive stabilization of membership in the skill-modulated network (fig 4a).

**Fig. 4:**
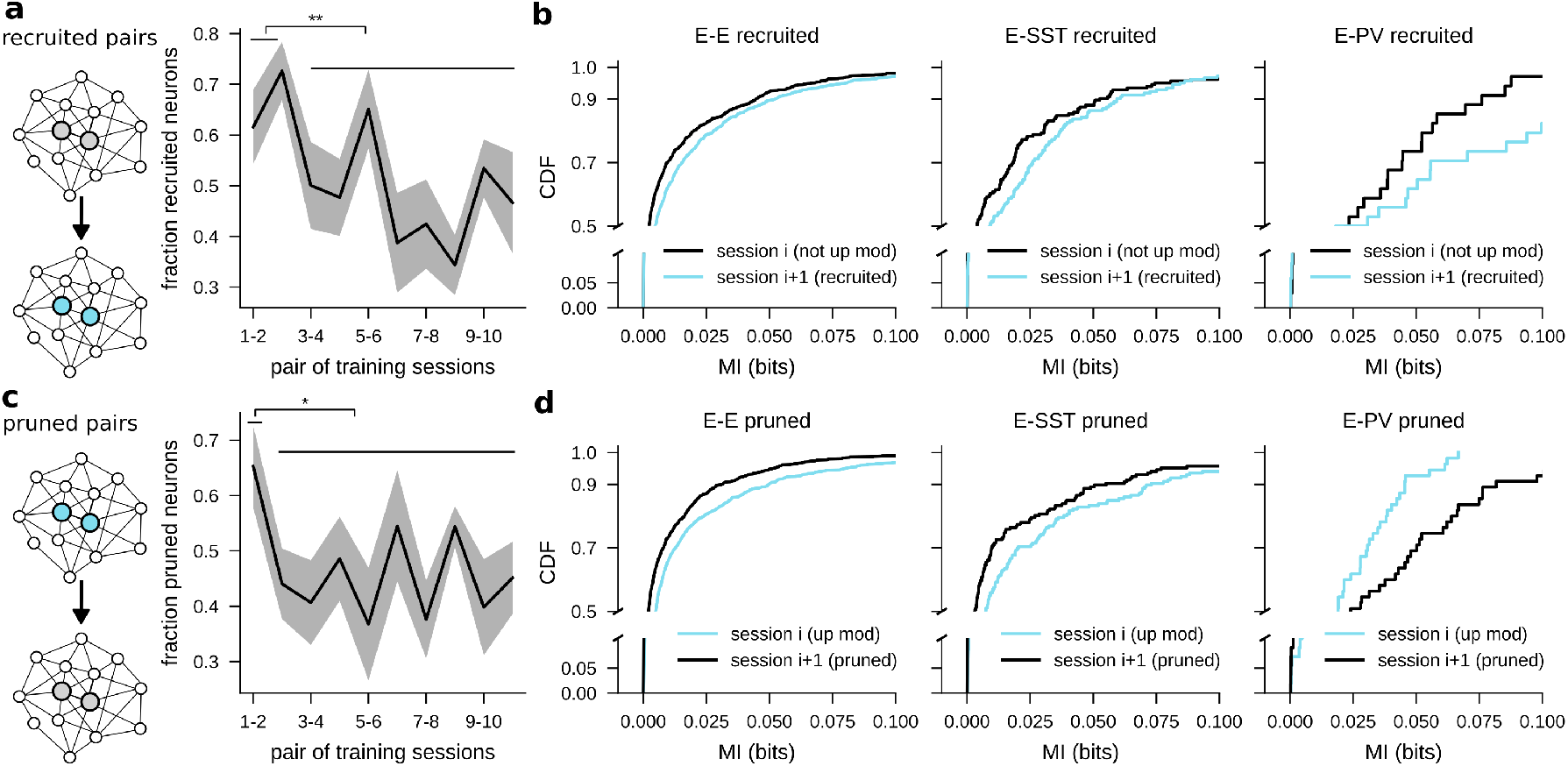
Changes in functional interactions track skill-modulated network recruitment and pruning. **a.** Left: schematic describing recruited pairs, in which both neurons in the pair transition into the skill-modulated network. Right: the fraction of neurons skill-modulated on a given session that were not skill-modulated on the previous session, for all adjacent session pairs. The fractions on session pairs 1-2 and 2-3 (n = 14 session pairs across 7 mice) are significantly higher than those on subsequent days (n = 56 session pairs across 7 mice). Mann-Whitney U-test p = 0.005, rank biserial r = 0.49. **b.** Empirical CDFs of MI for recruited pairs before and after recruitment. Only pairs with significant MI on both sessions are compared. *** indicates *p <* 0.001 on Wilcoxon signed-rank test of MI on session *i* + 1 vs MI on session *i*. Cyan indicates MI on the session in which the pair was skill-modulated. Left: E-E (n = 1100 neuron pairs, rank biserial r = 0.35). Middle: E-SST (n = 177 neuron pairs, rank biserial r = 0.38). Right: E-PV (n = 34 neuron pairs, p = 0.055). Again, cyan indicates MI on the session in which the pair was skill-modulated. **c.** Left: schematic describing pruned pairs, in which both neurons in the pair transition out of network membership. Right: the fraction of neurons skill-modulated in a given session that are no longer skill-modulated in the next session, for all adjacent session pairs. The fractions on session pairs 1-2 (n = 7 pairs across 7 mice) are significantly higher than those on subsequent days (n = 63 session pairs across 7 mice). Mann-Whitney U-test p = 0.025, rank biserial r = 0.52. **d.** Empirical CDFs of MI for pruned pairs before and after pruning. Only pairs with significant MI on both sessions are compared. *** indicates *p <* 0.001 on Wilcoxon signed-rank test of MI on session *i* vs MI on session *i* + 1. Left: E-E (n = 1040 neuron pairs, rank biserial r = 0.48). Middle: E-SST (n = 174 neuron pairs, rank biserial r = 0.46). Right: E-PV (n = 55 neuron pairs, p = 0.08).

High recruitment was most evident in the first four sessions (fig 4a). When both neurons in a pair were recruited into the skill modulated network, their MI increased across all cell types (fig 4b). Conversely, when both neurons in a pair lost skill modulation, MI decreased (fig 4d). This pruning of skill-modulated cells was most common from the first to second sessions, but continued throughout learning (fig 4c). Thus, changes in skill modulation were accompanied by coordinated changes in pairwise interaction strength: recruitment into the skill-modulated network involved not simply greater activation but also stronger pairwise temporal coordination, whereas loss of skill modulation was associated with weaker coordination. When cells are recruited to skill-modulation, they join a temporally coordinated and spatially specific pattern of activity.

### SST-associated coordination extends beyond the skill-modulated network and stabilizes with learning

Because skill-modulated neurons comprised less than half of the recorded population, we next asked whether structured interactions also extended to neurons that were not classified as skill-modulated. Importantly, skill modulation is a relative measure: it identifies neurons whose activity is greater during the skill epoch than during other behavioral periods, but does not capture the full temporal structure of their task-related activity. We therefore used multiresolution consensus clustering to identify groups of neurons with coherent interaction structure independent of their skill-modulation classification [27]. This approach applies Louvain clustering across a range of resolution parameters and randomized initializations, then identifies communities that are representative of the ensemble of partitions.

The resulting clusters contained mixtures of skill-modulated and non-modulated neurons, indicating that functional organization extended beyond the boundaries of the skill-modulated ensemble (fig 5a). Within clusters, SST-excitatory and PV-excitatory pairs exhibited greater MI than putative excitatory-excitatory pairs, recapitulating the preferential coordination observed within the skill-modulated ensemble (fig 5b). Cluster membership became more stable across sessions with learning (fig 5c), as measured by adjusted mutual information between cluster structure [28], which measures the similarity of cluster assignment adjusted for chance based on the size and number of the clusters. This increase in stability is consistent with progressive stabilization of the broader interaction structure. Thus, strong SST-excitatory interactions were not restricted to neurons identified by skill modulation, but were embedded within broader functional communities whose organization became increasingly stable over training.

**Fig. 5:**
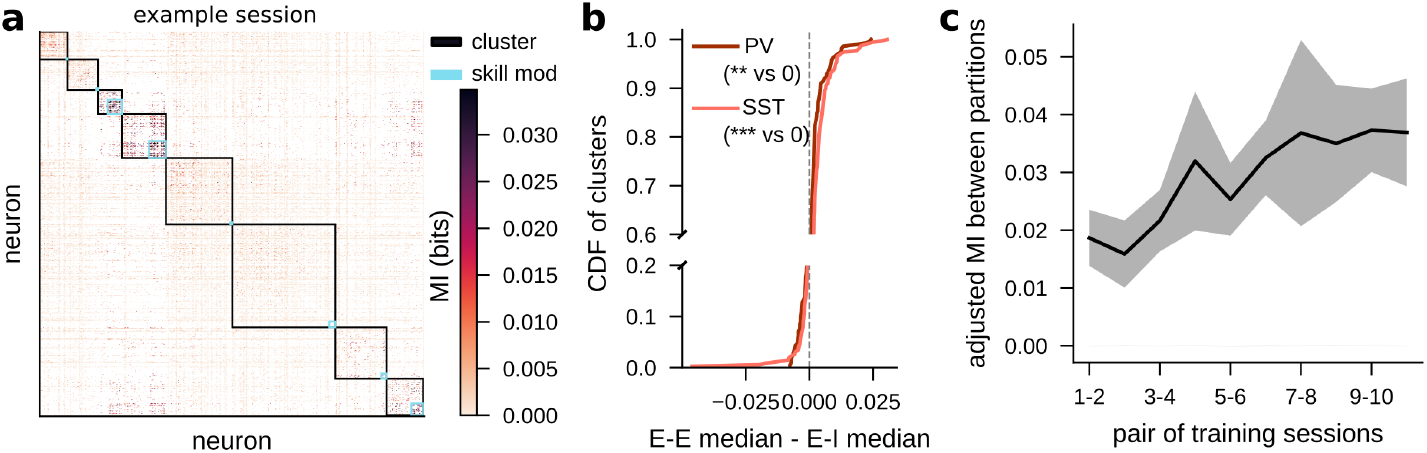
SST-associated coordination extends beyond the skill-modulated network and stabilizes with learning. **a.** Example session MI heatmap, sorted by cluster assignment. Skill modulated neurons in each cluster are designated by cyan boxes. **b.** Empirical CDF of the difference between the median of E-E edges and the median of E-I edges per cluster, for PV and SST. ** indicates *p <* 0.01 and *** indicates *p <* 0.001 in a Wilcoxon signed rank test against 0. **c.** Adjusted mutual information between cluster partitions on each pair of training sessions. SEM over mice (n=7).

### SST neurons and highly connected excitatory neurons carry more behavioral information than randomly selected excitatory neurons

To measure whether the information neurons carried about behavior differed based on cell type and coupling, we compared ridge regression performance for models trained to predict 3D reconstructed kinematics (fig 6a) from the activity of three different neuronal subsets. We compared inhibitory neurons to size-matched subsets of excitatory neurons defined based on the total MI (i.e. weighted degree) in their functional interactions. The highly connected subset included the neurons that were most strongly coupled to the rest of the population, as measured by the sum of their significant mutual information with all other cells (fig 6b). To select neurons that were strongly integrated with both excitatory and inhibitory neurons, we ranked neurons by the product of their z-scored excitatory and inhibitory functional interaction weights and selected top ranked neurons, matching the number of inhibitory neurons recorded in each session (fig 6b). This selected neurons that were strongly integrated into the local network. Because the total strength of inhibitory and excitatory functional connections were more correlated for SST neurons than for PV neurons (fig 2i), the highly coupled subset for SST mice included neurons that were particularly coupled with SST neurons. The third subset was composed of a random selection of excitatory neurons that did not fall into the highly connected subset, which typically included neurons whose coupling product was less than the 90th percentile (91.6 *±* 2.2 % of excitatory neurons).

**Fig. 6:**
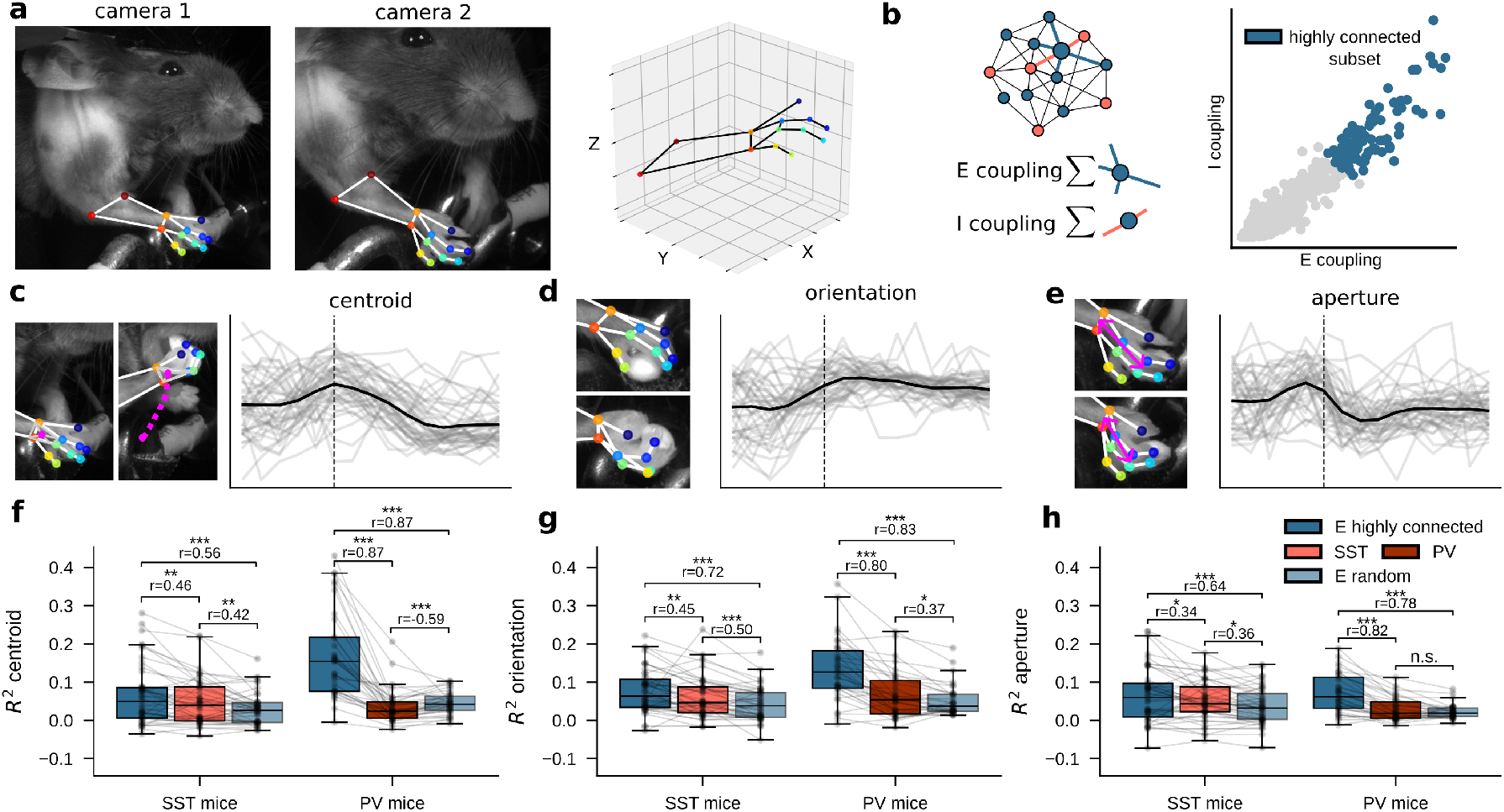
SST neurons and highly connected excitatory neurons carry more behavioral information than randomly selected excitatory neurons. **a.** Left, middle: two camera angles of behavioral video, with DeepLabCut keypoints and skeleton overlaid. Right: 3D triangulation of DeepLabCut keypoints. **b.** Schematic showing E and I coupling calculation based on total mutual information (i.e. weighted degree), and selection of the highly connected subset of excitatory neurons. **c.** Left: schematic describing paw centroid measure. Right: centroid traces for successful trials in an example session (gray) with mean overlaid (black). Dashed line shows pellet contact. **d–e.** Same as c for orientation and aperture. **f.** Ridge regression performance for models trained on highly connected excitatory neurons, SST or PV neurons, and randomly selected excitatory neurons. Individual sessions overlaid; n = 45 sessions for SST mice and n = 34 sessions for PV mice. *** indicates *p <* 0.001, ** indicates *p <* 0.01, and * indicates *p <* 0.05 on paired Wilcoxon signed-rank test, and r indicates rank biserial correlation. **g–h.** Same as f. for orientation and aperture.

We compared performance for models trained to predict three kinematic summary variables that we calculated from 3D reconstruction of paw and digit keypoints derived from two different camera angles (fig 6a). The paw centroid (fig 6c) quantifies the gross movement of the paw, the orientation (fig 6d) quantifies the twisting movement animals make as they grasp the pellet, and the aperture quantifies the motion of opening the paw and then closing it around the pellet (fig 6e). We compared model performances within the same sessions so that kinematics and trial counts were identical.

Models trained on highly connected excitatory neurons consistently out-performed models trained on the other neuronal subsets (fig 6f-h). Models trained on SST neurons also consistently outperformed models trained on randomly selected neurons (fig 6f-h). In contrast, models trained on PV neurons showed no consistent advantage over random excitatory subsets, performing sometimes worse (fig 6f), sometimes slightly better (fig 6g), and sometimes equivalently (fig 6h). Thus, excitatory neurons that are strongly integrated into the functional network carry more information about movement kinematics than weakly integrated neurons, and SST neurons carry more kinematic information than comparably sized random excitatory populations. Together with SST-excitatory preferential functional connectivity, these decoding results identify SST neurons and highly connected excitatory neurons as a behaviorally relevant functional ensemble whose activity carries information about movement kinematics.

## Discussion

With motor learning, SST activity becomes increasingly temporally consistent and coordinated with excitatory neurons in an emerging skill-related network. SST neurons were strongly recruited during skilled behavior and exhibited more reliable trial-related activity than either PV or putative excitatory neurons. This consistency emerged early in learning and correlated with behavioral improvement even after controlling for training session. Later in learning, strong coordination with a restricted subset of skill-modulated excitatory neurons emerged, with interactions dominated by aligned rather than opposing activity states. This coordination was dynamic across sessions: interaction strength increased as neurons were recruited into the skill-modulated network and decreased when skill modulation was lost. SST-associated organization also extended beyond the skill-modulated network, with SST neurons showing elevated coordination within MI-based functional communities containing both modulated and non-modulated neurons. Notably, SST neurons carried substantially more information about both paw position and paw posture than randomly selected sets of excitatory neurons. Together, these findings suggest that motor learning refines not only SST activity, but also which excitatory neurons SST cells coordinate with and when those interactions occur relative to behavior. Learning therefore refines the timing of SST inhibition, which supports the development of consistent dynamics in the excitatory population.

These results clarify conflicting findings in the field. Motor learning increases the consistency of M1 activity while reorganizing inhibitory circuits [1, 3], and SST axonal bouton density decreases with learning, whereas PV bouton density transiently increases [3]. Over a similar period, SST population activity increases [7]. Our results help reconcile increased SST activity with reduced SST bouton density by showing that learning is accompanied by stronger coordination between SST neurons and a sparse subset of skill-modulated excitatory neurons. Rather than reflecting a uniform increase or decrease in SST engagement, these findings suggest that learning increases the cellular and temporal specificity of SST-associated functional interactions.

By computing mutual information on concatenated single trial data, we quantified pairwise covariation within single trials rather than covariation between trial-averaged responses. Consistent trial-averaged activity was not required for strong functional interactions: despite low PETH consistency throughout training, PV neurons showed elevated MI with excitatory cells during early training. Using a trial-resolved measure of coordination is particularly important for variable behaviors such as reach-grasp-carry, in which both neural activity and behavior vary substantially across trials and trial-to-trial neural variability itself carries behaviorally relevant information [29, 30].We further isolated temporally specific interactions by comparing observed MI with a trial-rate-matched null. Together, this approach revealed that the emerging SST-excitatory interactions reflect reliable, temporally constrained alignment within single trials rather than similarity in trial-averaged responses. As M1 engaged in active, online control that varied moment to moment and trial to trial, these select SST-excitatory pairs were reliably recruited together, suggesting an activity motif that stabilized with learning. PV neurons showed significantly higher within and across session variability than SST neurons and excitatory neurons, despite being strongly recruited during the task. Their strong alignment with excitatory neurons early in training, even while their trial to trial consistency was low, implies that their activity closely tracked that of select excitatory neurons rather than the behavior. PV neurons were most strongly coordinated with excitatory neurons during the period when skill-related activity and the fraction of skill-modulated neurons were increasing most rapidly. This pattern is consistent with a role for PV recruitment in constraining or stabilizing the early expansion of skill-related cortical activity, given the transient increase in PV boutons during motor learning and the strong recurrent coupling between pyramidal and PV populations [3, 31, 32].

Together, these findings imply that PV and SST neurons contribute differently across phases of learning. PV-excitatory interactions were most prominent during early training sessions, whereas strong SST-excitatory interactions emerged as performance stabilized. SST neurons also participated during early learning: SST temporal consistency emerged early and covaried with behavioral improvement. Although turnover in the skill-modulated network neurons was highest early in training, network membership remained dynamic throughout behavioral refinement, and functional interactions changed accordingly. Interaction strength increased as neurons were recruited into the network, and decreased when skill-modulation was lost. This persistent coupling between network membership and interaction strength is consistent with SST-associated interactions contributing to the refinement and maintenance of activity patterns supporting skilled movement. Overall, the data support a phase-dependent shift in the relative prominence of PV- and SST-associated interactions.

This progression may be particularly relevant for behaviors that remain dependent on M1 after initial acquisition. Dexterous reach-to-grasp requires continued control of both gross transport and fine paw and digit configuration, and these components can improve over different timescales, with fine kinematics continuing to evolve after broader measures of performance have stabilized [16]. SST activity contained substantial information about both paw position and posture. These findings link the continued reorganization of SST-associated functional interactions to behavioral features that remain under refinement after initial task acquisition. These data support overlapping, phase-dependent contributions: early learning is accompanied by increasing temporal reliability of SST activity and strong PV-excitatory coordination, whereas later refinement is characterized by increasingly selective SST-excitatory coordination within the evolving skill-related population.

This organization extends beyond neurons classified as skill modulated. Because skill modulation reflects average differences in activity across behavioral epochs, neurons can have similar overall activity across behaviors while still showing temporally structured interactions within the behavior of interest. Functional communities defined independently of skill modulation contained both modulated and non-modulated neurons, became more stable with training, and exhibited preferentially strong SST-excitatory interactions. The behavioral relevance of this interaction structure was further supported by the decoding analysis. SST neurons and excitatory neurons strongly coupled to both SST and excitatory populations carried more information about movement kinematics than randomly selected excitatory neurons. Moreover, excitatory neurons with stronger excitatory functional connections also tended to have stronger SST connections, indicating that SST coupling was concentrated among highly integrated excitatory neurons. Thus, behavioral information is structured not only by the activity of individual neurons, but also by the functional interactions in which they participate. This extends previous evidence that dynamic M1 co-activity patterns carry behaviorally specific information [30, 33] by showing that such organization is cell-type specific and reorganizes with learning.

The changes observed here differ from those reported during visual associative learning, in which PV neurons become increasingly coupled to similarly selective pyramidal neurons, while SST neurons become decorrelated from the surrounding population [6]. This contrast argues against fixed computational roles for inhibitory subtypes across cortical areas or behaviors. In sensory associative tasks, behavior often depends on integrating population activity until stimulus identity can be classified, whereas motor control requires continuously coordinating activity to guide and refine movement in real time. SST and PV contributions may therefore depend on the computations required of the local circuit. Task dependence is also apparent within motor cortex. M1 is required for acquisition of all motor behaviors but can become dispensable for their execution after extensive training. For example, a learned, non-dexterous lever-press sequence can ultimately be executed without M1 [8, 9]. SST engagement similarly depends on task demands: SST activity becomes action locked during acquisition of a simple lever press but diminishes with extended training, whereas more temporally constrained sequences maintain or reorganize SST recruitment [10]. Together, these findings suggest that continued SST engagement is sustained when behavior continues to require precise temporal organization and refinement. More broadly, the contributions of SST and PV neurons appear to depend jointly on task demands and learning stage.

Overall, our results show that inhibitory contributions to motor learning are organized by cell type, neuronal features, and learning stage rather than by a uniform increase or decrease in inhibition across learning. PV-excitatory coordination is strongest during early learning, when skill-related activity is rapidly increasing, consistent with a role in constraining or stabilizing this initial reorganization. SST activity is also behaviorally relevant during early learning, but selective SST-excitatory coordination becomes increasingly prominent as learning proceeds. Critically, these functional interactions strengthen when neurons are recruited into the skill-related network and weaken when they leave it, directly linking functional interaction structure to the evolving composition of the cortical population corresponding to skilled behavior. These findings reveal a level of motor-cortical organization not captured by changes in neuronal activity alone: learning reorganizes which inhibitory and excitatory neurons interact, how strongly they interact, and when those interactions are engaged during skilled movement.

## Methods

### Mice

All experimental procedures were approved by the University of Chicago Institutional Animal Care and Use Committee and were performed in compliance with the Guide for the Care and Use of Laboratory Animals.

To label SST and PV neurons, we crossed mice expressing Cre-dependent td-tomato (B6.Cg-Gt(ROSA)26Sor(tm14(CAG-tdTomato)Hze/J, Jax strain 007914, commonly called Ai14) with either Sst-IRES-Cre (Ssttm2.1(cre)Zjh/J, Jax strain 013044, n=4) or PV-Cre (B6.129P2-Pvalbtm1(cre)Arbr/J, Jax strain 017320, n=3). Both male (n=4, 3 SST and 1 PV) and female (n=3, 2 PV and 1 SST) mice were used. Mice were between 10 and 11 weeks old at the start of the training protocol.

### Surgical Procedures

Mice underwent two surgeries: viral injection followed by cranial window and headplate implantation. For both procedures, anesthesia was induced with 3% isoflurane and maintained at 1–1.5%.

For viral expression of GCaMP6f, three burr holes were made over the caudal fore-limb area of primary motor cortex. AAV9 carrying GCaMP6f under the synapsin promoter (pAAV.Syn.GCaMP6f.WPRE.SV40; Addgene #100837-AAV9) [34] was injected at three stereotaxic locations relative to bregma: +0.25 mm anterior, 1.25 mm lateral; +0.25 mm anterior, 1.75 mm lateral; and 0 mm anterior, 0.5 mm lateral. These were designed to target caudal forelimb area [35]. At each site, 150 nL of undiluted virus was injected at depths of approximately 500 and 250 *µ*m below the cortical surface at 1 nL/s. The micropipette was left in place for at least 5 min after each injection before withdrawal. Mice recovered for at least 7 days before cranial window implantation.

For the second surgery, a custom titanium headplate was secured to the skull with dental cement. A 3.5-mm craniotomy centered over the injection sites was opened and the underlying dura removed. A 3.5-mm-diameter, 0.66-mm-thick glass plug bonded to a 5-mm coverslip was positioned within the craniotomy such that the plug was flush with the cortical surface. The outer coverslip rested on the surrounding skull and was secured with dental cement. Mice recovered for at least 7 days before habituation and behavioral training.

### Behavioral training

Mice were food restricted to approximately 90% of baseline body weight; animals falling below 85% were returned to ad libitum feeding until weight recovered. Before training, mice were habituated to handling and head fixation for approximately 1 week.

Training began with a shaping phase in which sucrose pellets were presented within licking distance. Once mice reliably retrieved pellets, pellets were placed in a recessed trough that required a forelimb reach. After at least 15 successful reaches from the trough, mice progressed to the pedestal-based task. This intermediate shaping step provided early successful trials before introduction of the more difficult pedestal configuration, similar to previous reach-training procedures [36].

Mice were positioned to promote use of the right forepaw, contralateral to the imaged hemisphere. If necessary, the left paw rest was raised to prevent use of the ipsilateral paw. Seating and head-fixation positions were adjusted individually to permit comfortable, unobstructed reaching. Pedestal position was optimized for each mouse and then held constant throughout training.

During training, 14 mg sucrose pellets (Bio-Serv) were presented individually on a rotating pedestal apparatus. A new pellet was delivered after the previous pellet was consumed, displaced from the pedestal, or left without a reach attempt for 30 s. Mice trained for 30 min per day at approximately the same time each day.

### Pellet Delivery Apparatus

Pellets were delivered using a custom apparatus consisting of a pellet dispenser (Campden Instruments), a 3D-printed funnel, and a 3D-printed rotating pedestal assembly driven by a servo motor and controlled using custom Arduino code. Design files and additional construction details are available upon request.

### Behavioral video and behavioral analysis

#### Video and keypoint tracking with DeepLabCut

Behavior was recorded throughout training at 200 frames/s using two offset cameras (FLIR Blackfly S BFS-U3-16S2C). One camera viewed the mouse approximately 30*^◦^* from the frontal axis and 20*^◦^* from above; the second viewed the mouse approximately 80*^◦^*from the frontal axis and near the horizontal plane. Both cameras were positioned to the mouse’s right. Paw and digit keypoints were tracked using DeepLabCut [37], including wrist, metacarpophalangeal, and proximal interphalangeal landmarks.

#### Trial segmentation

Candidate behavioral events were identified from peaks in the vertical position of the inner-wrist key-point from the more lateral camera using scipy function *`find_peaks`*. A subset of candidate events was manually annotated as reach, grasp, carry, fidget, grooming, food manipulation, or non-movement. These annotations were used to train a random-forest classifier (sklearn’s *`RandomForestClassifier`* class) using the 2D keypoints from both camera views. Classifier accuracy exceeded 80% on held-out data, after which the trained model was applied to the remaining candidate events. Automated classifications were additionally spot checked by experimenters for quality control.

We note that reach, grasp, and carry behaviors did not necessarily occur in sequence, since mice often performed several reaches before attempting to grasp, and several grasps before proceeding to a successful or empty carry. Accordingly, analyses of skill-related activity included all time points during which mice were actively engaged in reach, grasp, or carry behavior, including events occurring outside a complete reach-grasp-carry sequence.

#### Behavioral Success

To quantify success rate, we manually reviewed and classified behavior video surrounding grasp and carry behavioral events. We calculated success rate as the number of pellets successfully obtained divided by the number of pellets presented. The time of each pellet delivery was recorded with a short voltage pulse from the pellet delivery system to the integrated Bruker recording setup. Occasionally the pellet turntable was triggered but failed to actually deliver a pellet, so we used the DeepLabCut pellet keypoint to filter for delivery voltage pulses associated with a real pellet delivery. A pellet was determined to be present if the pellet likelihood was *>* 0.9 in keypoints from both camera angles for at least 125 ms of the timespan from 50 ms preceding to 200 ms following the pedestal stopping in front of the mouse. This amount of time was chosen after observing videos to account for spontaneous pellet dislodgement but not animal-related task behavior.

#### Quantification of improvements in success rate

We quantified improvements in success rate by performing logistic regression on binary trial outcomes vs session number using the sklearn *LogisticRegression* class for each mouse.

#### 3D triangulation

Two-dimensional keypoint trajectories were low-pass filtered using a zero-phase, 10th-order Butter-worth filter with a 15-Hz cutoff (scipy.signal.filtfilt). Samples with DeepLabCut likelihood *<* 0.4 were excluded. Missing segments of up to 20 consecutive frames (100 ms) were linearly interpolated; longer gaps were left missing. Three-dimensional coordinates were then reconstructed from the two camera views using the DeepLabCut 3D pipeline [38].

#### Kinematic variable calculation

Three kinematic variables were calculated using 3D triangulation data: paw centroid, orientation, and aperture. Paw position was defined as the centroid of four landmarks: the digit 2 and digit 3 metacarpophalangeal joints and the radial and ulnar wrist landmarks. Aperture was defined as the distance between the midpoint of the digit 2 and digit 3 proximal interphalangeal joints and the midpoint of the radial and ulnar wrist landmarks. Digit-tip landmarks were excluded because they were frequently occluded. Paw orientation was estimated from the normal vector to the plane defined by the digit 2 metacarpophalangeal joint, digit 3 metacarpophalangeal joint, and outer-wrist landmark. The *y*-component (as in fig 7a) of this normal vector was used as the orientation measure, with the *y*-position of the wrist subtracted so that negative values indicated palm-down orientations and positive values indicated palm-up orientations.

#### Two-photon calcium imaging

Two-photon calcium imaging was performed using a Bruker microscope in a single imaging plane at 30 Hz. We imaged at approximately 200–300 microns below the surface. The field of view (FOV) was 828 *×* 828 microns, sampled at 512 *×* 512 pixels. At the beginning of each session, tdTomato fluorescence from SST or PV interneurons was recorded for approximately 2 min using 1040-nm excitation. GCamp6f fluorescence was then recorded throughout the rest of the session with a laser wavelength of 920 nm.

To maintain a consistent field of view across sessions, imaging location was first aligned using the surface vasculature. The red-channel image was then matched to the previous session by adjusting *x*-*y* position and sampling focal planes at the prior imaging depth and at *±*5, *±*10, and *±*15 *µ*m. The plane providing the closest match to the previous session was selected for imaging.

### Calcium imaging preprocessing

#### Region Of Interest (ROI) extraction and dF/F calculation

ROIs were extracted using Suite2p (v0.14.5) [39] from the raw images. ROIs were then classified as cells or non-cells based on a classifier trained on several manually curated FOVs. After applying the classifier, we additionally manually curated the data to ensure accuracy. For each cell, neuropil fluorescence was subtracted using a cell-specific coefficient selected to optimize trace smoothness, based on a modification of the Allen Institute optical physiology pipeline (https://github.com/AllenInstitute/ophys_etl_pipelines/tree/main). Δ*F/F* was then calculated using a median-filter-based estimate of baseline fluorescence adapted from the same pipeline.

#### Spike inference

Inferred spiking activity was estimated from Δ*F/F* traces using the CASCADE algorithm [40], using the `Global_EXC_30Hz_smoothing50ms_high_noise` models. CASCADE selects among models matched to the noise level of each neuron, estimated from frame-to-frame fluorescence fluctuations. The same model family was applied to excitatory, SST, and PV neurons because inhibitory-specific GCaMP6 models did not provide improved performance at the time of analysis. CASCADE outputs the expected number of spikes per imaging frame; these values were multiplied by the 30-Hz imaging rate to obtain inferred firing rates in spikes/s.

#### Registering cells across sessions

Cells were registered across sessions using CellReg [41]. For our analyses, we focused on pairs of adjacent sessions to maximize the number of registered cells.

#### Statistical tests

We used the Wilcoxon rank sum test to evaluate the significance of results in paired data, including MI for the same neuron pair on different days and decoding performance from different neural subsets on the same kinematic data. We used the Mann-Whitney U-test for unpaired data, including MI for neuron pairs including different cell types. We chose these tests because they do not assume normality. To measure trends over training sessions, we used linear mixed effects models fit to the data, with a Wald test on the fixed effect coefficients. To quantify increases in the success rate over training, we fit a logistic regression on binary trial outcome per mouse. We chose this approach, instead of simply fitting a model to success rates, to account for the number of attempts.

#### Population activity and modulation depth

We calculated population activity by averaging inferred firing rate over trials and over neurons of each cell type. We calculated modulation depth of the population activity as the maximum population activity in the trial window divided by the median of population activity during classified, non-skill timepoints (fidget, food manipulation, non-movement, and grooming).

#### Peri-event time histograms (PETHs)

We calculated each neuron’s peri-event time histogram, or PETH, as the average of its inferred firing rate from 167 ms (5 imaging bins) before behavior start to 333 ms (10 imaging bins) after behavior start.

#### Split Session Correlations

Within each session, we took 1,000 random draws of approximately half of all skill trials. For each split, peri-event time histograms (PETHs) were calculated separately from the two subsets, and their similarity was quantified using Pearson correlation. Correlation coefficients were Fisher *z*-transformed before averaging across the 1,000 splits for each neuron. Neuron-level values were then averaged across cells in Fisher *z*-space and back-transformed for visualization.

Learning related changes in split session reliability were quantified using a linear mixed effects model (MixedLM, *statsmodels*) with session number and trial count as fixed effects and mouse as a random intercept. Trial count was included as an additional covariate in the model, since trial number increased with session number and could have contributed to better estimated PETHs. Statistical analyses were performed on Fisher *z*-transformed correlation values.

#### Partial correlation between split half correlation and success rate

We computed the partial correlation between the mean split half correlation on each session and the session’s success rate, controlling for session number, since both measures increased with session number. We first performed linear regression using scipy’s *linregress* function between correlation and session number, and between success rate and session number, and then calculated the Pearson correlation between the residuals.

#### PETH correlations across sessions

Across-session reliability was measured for neurons registered by CellReg on each pair of adjacent sessions. PETHs were computed separately for each component of skilled behavior (reach, grasp, and carry), and the Pearson correlation was computed between the PETH for each component on each session. The three component-specific correlations were Fisher *z*-transformed, averaged, and back-transformed to obtain a single across-session reliability value for each neuron and adjacent-session pair.

#### Skill modulation

We calculated an ANOVA for each neuron between activity during skill trials and activity during classified, non-skill trials (fidget, food manipulation, non-movement, and grooming). We first calculated the mean of timepoints within each trial by locking to behavior start and computing the average with the same time bins as the PETH calculations. This made the measurements in the distribution more independent by accounting for the autocorrelation in adjacent timepoints. A neuron was considered modulated if it had *p <* 0.05*/n*, where *n* is the number of neurons in the session.

### Mutual information

#### Adaptive Binning

We adaptively binned inferred spike counts from CASCADE. We first thresholded the inferred spike counts at a fixed threshold of 0.05, and placed all values below this threshold in the first bin. We then binned the remaining values such that each bin received an equal number of points across the entire session. To achieve this, we calculated the cumulative distribution function of the log-transformed spike count, and divided it into five equal bins. We then reverse transformed these bins to get the edges of each bin in spike count space.

#### MI calculation

We calculated the mutual information between the binned activity of each pair of neurons as:

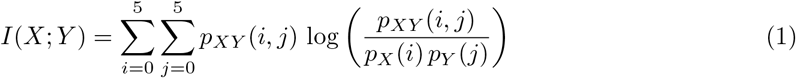

We used concatenated activity from every skill trial, using the same behavior locking and times per trial as the PETH calculations.

#### Trial-rate matched shuffle null

We shuffled activity and recalculated MI for each pair of neurons 1000 times. The shuffle was performed per trial by circularly shifting each trial’s 15 time bins by 3 to 10 bins, where the number of bins to shift was selected randomly per trial. To calculate significance for each neuron pair, we computed empirical *p* values as the fraction of times the MI for the shuffled activity exceeded or equaled the true MI. A neuron pair was considered significant if it had *p <* 0.05 after Benjamini-Hochberg false discovery rate correction.

#### Pointwise MI

We decomposed MI into its pointwise components, where

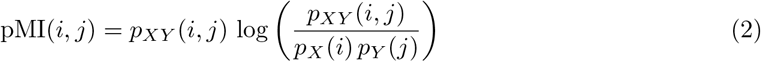

[17]. Because we used 6 bins, there were 36 pointwise components. We defined the subset of components with aligned states as the components where: (*i* = 0*, j* = 0), (*i* = 1*, j* = 1), (*i* = 2*, j* = 2), (*i* = 3*, j* = 3), (*i* = 4*, j* = 4), (*i* = 5*, j* = 5). Similarly, we defined the subset of components with anti-aligned states as the components where: (*i* = 0*, j* = 5), (*i* = 0*, j* = 4), (*i* = 1*, j* = 5), (*i* = 5*, j* = 0), (*i* = 4*, j* = 0), (*i* = 5*, j* = 1). We summed over these components to quantify the aligned and anti-aligned contributions to the total MI. For a similar method, see Ghirga et al., 2021 [18].

#### Clustering

We used multi-resolution consensus clustering [27] to identify clusters in the MI. The implementation uses functions from the Brain Connectivity Toolbox in MATLAB [42] and is identical to the implementation in Pope et al., 2021 [43].

### Ridge regression models

#### Interpolation

Kinematic variables sampled at 200 Hz were interpolated to the 30-Hz neural imaging timestamps using numpy.*interp*.

#### Functional coupling calculation and neuronal subset selection

For each excitatory neuron, excitatory and inhibitory functional coupling were defined as the summed MI of its significant interactions with excitatory and inhibitory neurons, respectively. To compare neuronal populations of equal size, excitatory subsets were matched to the number of labeled inhibitory neurons recorded in each session.

Highly connected excitatory neurons were selected based on their joint coupling to excitatory and inhibitory populations. Excitatory and inhibitory coupling were normalized separately across excitatory neurons to account for the substantially different numbers of potential excitatory and inhibitory partners, and neurons were ranked according to the product of their excitatory and inhibitory coupling. The top *n* neurons, where *n* equaled the number of labeled inhibitory neurons in that session, comprised the highly connected excitatory subset. A comparison subset of *n* excitatory neurons was sampled without replacement from the remaining excitatory population using numpy.random’s *`default_rng`* class.

#### Model fitting

Separate ridge regression models (sklearn’s *Ridge*) were trained to predict each kinematic variable: centroid, orientation, and aperture. Analyses included all timepoints from concatenated skill trials for which corresponding kinematic variables could be estimated. Data was divided into train and test sets using sklearn’s *`RepeatedStratifiedKFold`* class. Cross-validation was performed at the trial level to prevent temporally adjacent samples from the same trial from appearing in both training and test sets. Train/test splits were stratified based on behavioral component so that the distribution of reach, grasp and carry trials was similar in the train and test sets. Five-fold cross validation was repeated 1000 times for the highly connected excitatory and the inhibitory subsets. For the random excitatory comparison, we performed 100 random draws of excitatory neurons with 10 repeated five-fold splits each. On each train set, we performed an inner cross validation with five folds to estimate the optimal regularization parameter. Model performance was quantified with the coefficient of determination (*R*^2^, sklearn.metrics.*`r2_score`*) between true and predicted kinematics.

## Supplementary information

Supplementary figures are available in a separate document.

## Acknowledgements

We thank Elizabeth de Laittre, Harold Rockwell, Zachery Morrissey, Maria Pope, John McCann, Sunnie Hong, Stephanie Palmer, Nicholas Hatsopoulos, Brent Doiron, and Monica Rosenberg for helpful discussions and comments on the manuscript. We thank Elizabeth de Laittre and Harold Rockwell for technical assistance.

## Declarations

### • Ethics approval and consent to participate

Not applicable.

### • Consent for publication

Not applicable.

### • Data availability

Data is available upon request.

### • Materials availability

This study generated no new materials or reagents.

### • Code availability

Code used to generate the figures will be available on GitHub at time of publication.

### • Author contribution

G.W.F.: Conceptualization, Methodology, Software, Validation, Formal analysis, Investigation, Data curation, Writing – original draft, Writing – review & editing, Visualization, Funding acquisition.

E.T.W.: Software, Investigation, Data curation, Writing - review & editing.

J.N.M. Conceptualization, Methodology, Writing - original draft, Writing - review & editing, Supervision, Project administration, Funding acquisition.

