## Supplemental Information for "Somatostatin-excitatory coordination emerges with dexterous motor skill learning"

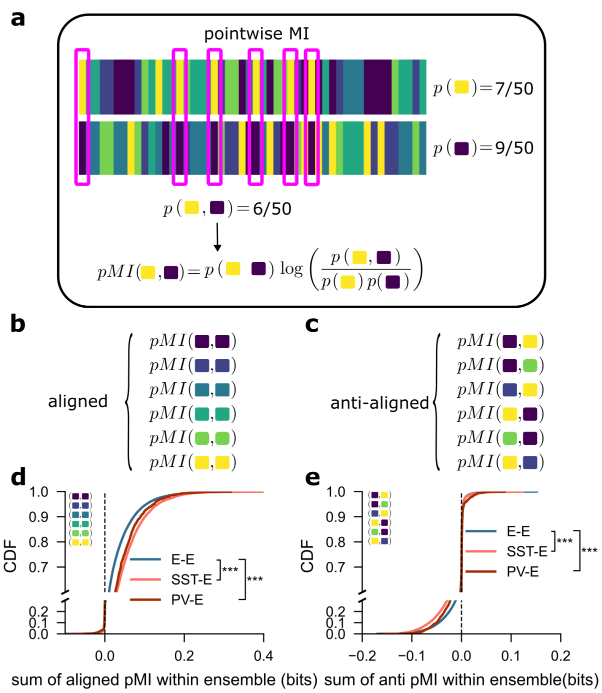


**Figure 1: Elevated mutual information reflects aligned rather than anti-aligned activity. a.** Schematic describing pointwise MI decomposition. **b–c.** Pointwise MI components are divided into aligned and anti-aligned groups. **d.** Empirical CDFs of the sum of aligned pMI components within the skill-modulated ensemble for E-E (n = 76079 pairs), E-SST (n = 12208 pairs), and E-PV (n= 2052). *** indicates $p < 0.001$ with Mann-Whitney U-test. SST-E vs E-E rank biserial r = 0.18, and PV-E vs E-E rank biserial r = 0.12. **e.** Same as d. for anti-aligned pMI components. *** indicates $p < 0.001$ with Mann-Whitney U-test. SST-E vs E-E rank biserial r = -0.17 and PV-E vs E-E rank biserial r = -0.15.

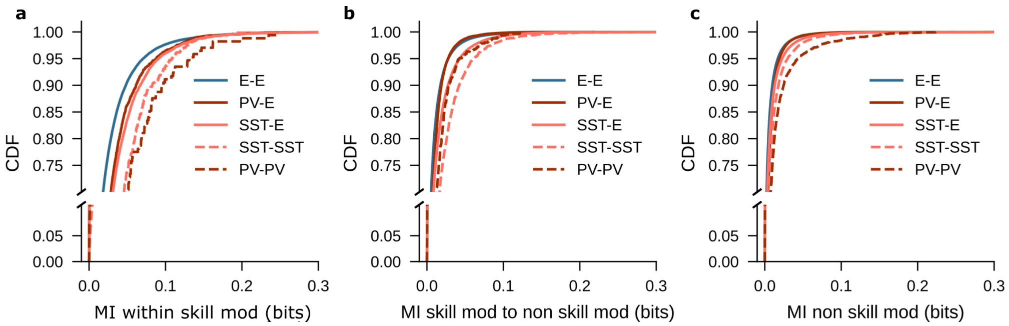


**Figure 2: MI is elevated in PV-PV and SST-SST pairs both within and outside the skill-modulated network. a.** MI between skill-modulated pairs, separated by cell type. **b.** MI between skill-modulated and non-skill-modulated neurons, separated by cell type. **c.** MI between non-skill-modulated pairs, separated by cell type.
